# BASIS: BioAnalysis SEDFIT Integrated Software for cGMP Analysis of SV-AUC Data

**DOI:** 10.1101/2023.10.12.562024

**Authors:** Alexander E. Yarawsky, Erik S. Gough, Valeria Zai-Rose, Natalya Figueroa, Hazel M. Cunningham, John W. Burgner, Michael T. DeLion, Lake N. Paul

**Author notes:** Authors contributed equally to the manuscript (Alexander E. Yarawsky, Erik S. Gough).

## Abstract

Sedimentation velocity analytical ultracentrifugation (SV-AUC) has long been an important method for characterization of antibody therapeutics. Recently, SV-AUC has experienced a wave of new interest and usage from the gene and cell therapy industry, where SV-AUC has proven itself to be the “gold-standard” analytical approach for determining capsid loading ratios for adeno-associate virus (AAV) and other viral vectors. While other more common approaches have existed in the realm of cGMP-compliant techniques for years, SV-AUC has long been used strictly for characterization, but not for release testing. This manuscript describes the challenges faced in bringing SV-AUC to a cGMP environment and describes a new program, “BASIS”, which allows for 21 CFR Part 11-compliant data handling and data analysis using the well-known and frequently cited SEDFIT analysis software.

## Introduction

Analytical ultracentrifugation (AUC) is a highly regarded first-principle technique with nearly 100 years of use in the characterization of biological materials ^1,2^. The technique is extremely versatile, being suitable to characterize macromolecules ranging from small peptides and organic compounds to antibodies and antibody-drug conjugates to the ribosome and chromatin complexes to viruses and lipid nanoparticles ^3–11^. With such a wide range of applications, it’s easy to imagine how useful AUC can be for characterization of biopharmaceuticals. Whereas antibodies have long been examined by sedimentation velocity AUC (SV-AUC) for aggregate quantitation ^4,5,10,12–16^, of particular interest recently is the use of SV-AUC for determination of loading states among adeno-associated virus (AAV) and other viral vectors ^8,17–24^.

Clearly, there is a critical need to facilitate the use of SV-AUC analyses in filings with regulatory agencies such as the US Food and Drug Administration (FDA) and the European Medicines Agency (EMA). However, the data collection and data analysis present numerous challenges that have – until recently – prevented simple and complete translation of the technique into the current Good Manufacturing Practices (cGMP) space. Previous software packages have circumvented one or more of these challenges. John Philo’s SVEDBERG, DCDT+, and kDalton programs ^16,25^ contain comprehensive logs that act as an audit trail. This suite of programs also allows the user to save a complete analysis within a single file, such that all necessary information is contained and available to readily replicate the analysis. This is in contrast to the current standalone SEDFIT that requires the raw data files and several different accessory files to be maintained in specified directories if the analysis is to be appropriately reloaded. Borries Demeler’s recent ULTRASCAN GMP module is designed to function under cGMP ^26^. The ULTRASCAN GMP module uses a designated server for its high-performance computing required for data analysis, and it also recommends the use of intensity measurements rather than absorbance measurements. The ULTRASCAN GMP module is currently under development to accommodate 21 CFR Part 11 guidance. In this manuscript, we describe a new program (“BASIS: BioAnalysis SEDFIT Integrated Software”) which contains and controls SEDFIT – the most widely cited AUC analysis software – via a completely cGMP-compliant environment that is 21 CFR Part 11 compliant and has undergone computer software validation.

### Typical Workflow for cGMP SV-AUC Analysis

The standard SV-AUC workflow includes several vulnerabilities and challenges for cGMP compliance (Figure 1) (See Savelyev, et al. ^26^ for a comprehensive discussion). The following observations are applicable to the Beckman Coulter Optima XL-A/XL-I instruments released in the early 1990’s, as well as the later Beckman Coulter Proteome Lab XL-A/XL-I instruments. Both instruments now use the same Proteome Lab data collection software. The newer Optima AUC released in 2017 by Beckman Coulter contains major changes to the instrument and control software. Specific comments will be made where the Optima AUC offers improved solutions over previous models, however, there are still vulnerabilities present with the newer Optima AUC software.

**Fig. 1.**
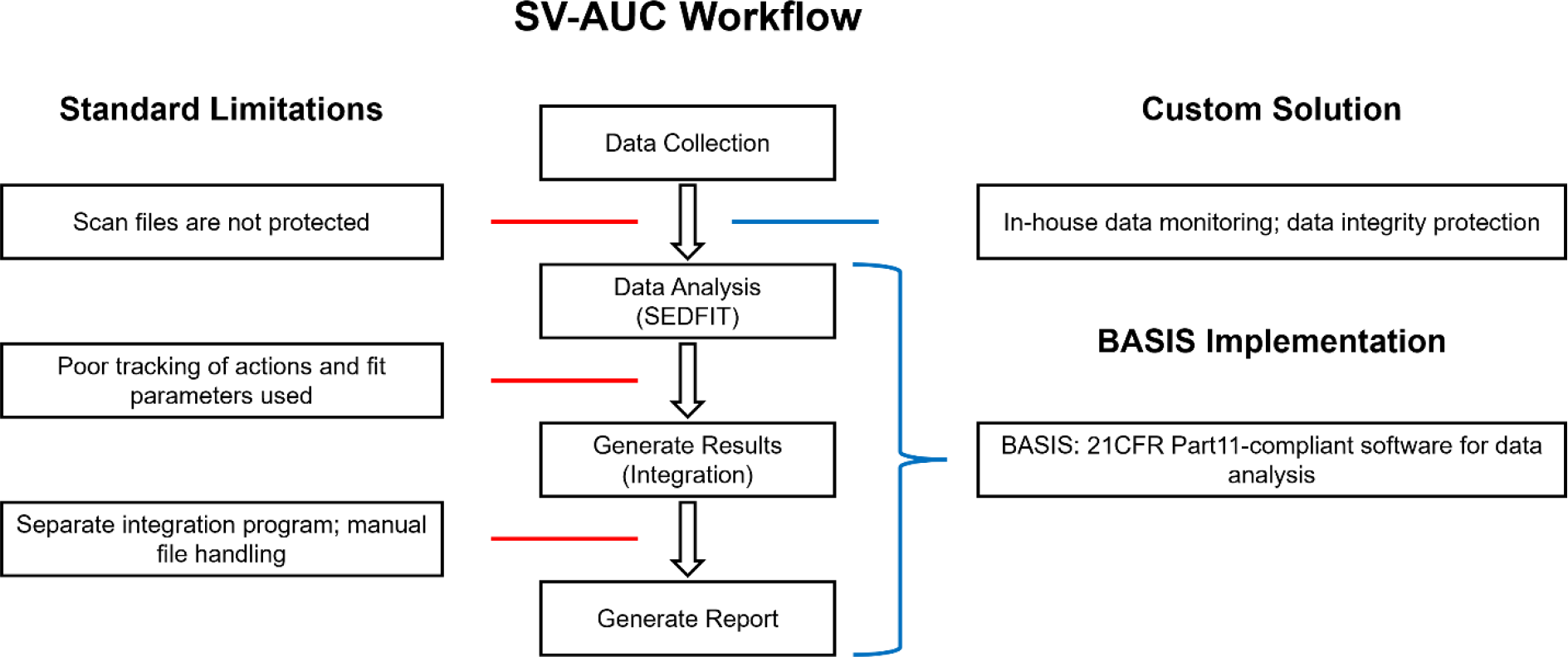
A typical SV-AUC workflow is presented. Limitations of each step relating to cGMP-compliance are noted to the left with red lines. The improvements upon each step to attain cGMP-compliance are shown to the right with blue lines.

Data collection exposes the first vulnerability in an SV-AUC workflow; in that, SV-AUC experiments involve collecting scans across the radius of a sample cell as the particles in solution sediment over several hours or overnight. The scans are then analyzed, usually by fitting to a model describing sedimentation and diffusion (i.e., the Lamm Equation ^27,28^), producing a distribution of species present within the sample. Data (ASCii) files are saved locally to the controlling computer after each scan is completed by the instrument. These files are immediately accessible to the user, and they are simple text files whose contents can be edited manually. The Optima AUC offers an improvement, as it compiles all data (ASCii files) into a compressed archive before making it available to the user. A simple solution to this first vulnerability on the XL-A/XL-I instruments involves monitoring the instrument’s controller (computer) for the generation of new files in the data directory, logging of the experiments, and implementation of signature requirements. Once a file is generated, it can be saved to a protected location where it cannot be altered. The experimental details and signatures can be mitigated with Standard Operating Procedures (SOPs). Scripts to examine the meta-data of the files or to compare them with the instrument’s generated log file will provide further assurance that data were not manipulated and are acceptable for analysis.

Data analysis presents the most complicated challenge to performing cGMP SV-AUC analysis. Several different analysis software are available, where the most common approach for biotherapeutic applications (e.g., antibody trace aggregate quantitation and AAV capsid loading) is the continuous c(*s*) distribution implemented in SEDFIT ^5,10,18–21,27^. SEDFIT analysis begins with loading of the data, selecting a model and fitting parameters, executing the fit, and exporting the final c(*s*) distribution. Until the recent release of SEDFIT v16.50 ^29^, external control of SEDFIT did not exist, nor could SEDFIT export complete information on many aspects of the executed fit. The FDA requires that any software used during a cGMP analysis be compliant with 21 CFR Part 11 from the Code of Federal Regulations. Until now, the best approach to attempt to meet this compliance was to apply SOPs, including copying screenshots of analyses.

This still omits important fit information, and more importantly, does not truly meet the compliance as it lacks various aspects of data integrity. Another vulnerability in the typical SV-AUC workflow involves transferring the calculated c(*s*) distribution from SEDFIT into a different program for peak integration and plotting (e.g., Excel, Origin, GUSSI, ChemStation, Empower, etc.). While SEDFIT does allow for peak integration within the program itself, this is not recommended, since the range often needs specifying manually with the cursor. There is also no way to export the resulting values, thus, they would need to be manually inserted into another program for calculation of averages, standard deviations, etc. It’s also typical that the area from more than one range would need to be examined for the complete result (i.e., mAb monomer vs dimer, AAV empty vs full), again indicating the need to perform the integration in a separate program.

### BASIS Workflow for 21 CFR Part 11 cGMP SV-AUC Analysis

The objective of the BASIS software involves condensing the Experimental Details, Data Analysis, Integration, and Reporting into a single, protective platform that is convenient and compliant with regulations for cGMP analysis. This software was made possible by the release of the command line-interactive SEDFIT v.16.50 ^29^. BASIS is a container for SEDFIT and several other components that allow for distribution integration, calculation, and report generation – all while maintaining an audit trail and reducing user intervention. Furthermore, the interface was developed with additional concerns in mind, which is to enhance the analyst’s ability to examine the raw data and evaluate the fit while allowing Quality Control (QC) review of the experiment and final data analysis. Figure 2 illustrates the general workflow for cGMP analysis of SV-AUC data using BASIS.

**Fig. 2.**
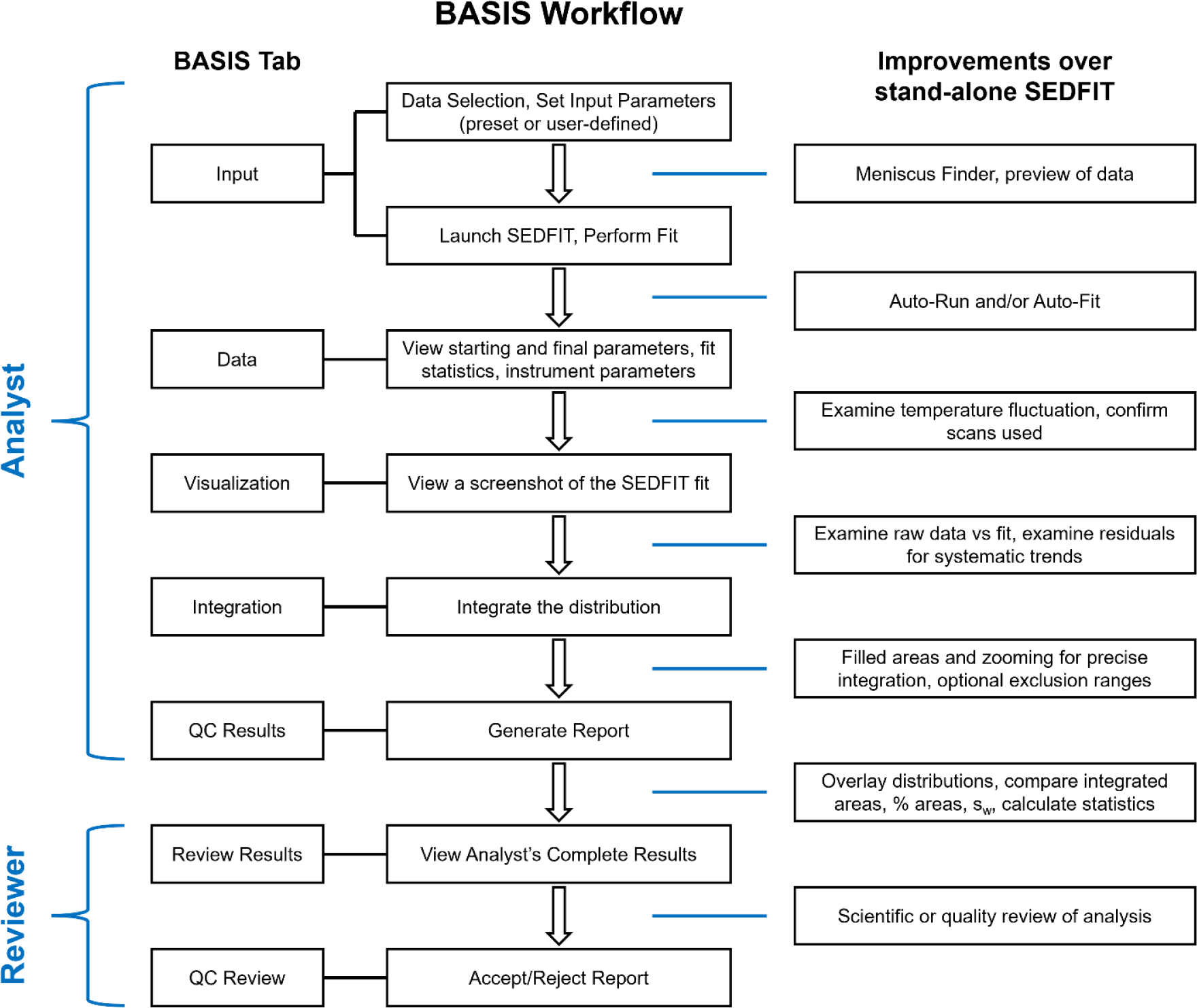
The BASIS workflow is shown as a flow chart. The center of the chart briefly describes each step. The left side of the chart (BASIS Tab) denotes the various tabs or pages within BASIS that relate to the step. The right side of the chart offers additional insights into the steps and how they are mediated by features available in BASIS.

### 1. Analyst: Input Tab

The first step in BASIS is to select a parameter set for analysis. BASIS offers the capability to save input parameters specific to different sample types. For example, an AAV sample requires a much wider range of sedimentation coefficients and a different starting frictional ratio (*f*/*f*_0_) than, for example, a mAb sample. Regardless of the Parameter Set chosen, all the initial and final fitting parameters are saved. BASIS also implements a meniscus-finding algorithm. This routine examines the raw data in a directory to find the meniscus, automatically sets the meniscus and meniscus fitting limits, the left fit limit of the useful data, and provides the analyst with a preview of the data. The analyst can readjust limits if needed during this preview. SEDFIT is then launched and an Auto-Run and/or Auto-Fit can be performed, or the analyst can operate SEDFIT traditionally within the SEDFIT window. Once the analysis is finished, SEDFIT is redrawn for a screenshot of the data fit range, with fit statistics shown (equivalent to the Ctrl+D and Ctrl+O keyboard commands). Finally, SEDFIT is closed, and new tabs appear in BASIS.

### 2. Analyst: Data Tab

The Data Tab in BASIS is divided into several sections. The first section includes the calculated c(*s*) distribution, the sample name, the SEDFIT version, and the data files used in the analysis. The next section lists various experimental parameters, such as rotor speed, wavelength, and temperature information. The remainder of the Data Tab displays the initial and fitted parameters of the model, along with the fit statistics. This tab is read-only – no information may be altered by the user.

### 3. Analyst: Visualization Tab

The Visualization Tab displays the screenshot of the SEDFIT window after the fitting is completed. This provides the third opportunity for the analyst to examine the raw data, as well as the second opportunity to evaluate the fit via the residuals, histogram plot, and c(*s*) distribution.

### 4. Analyst: Integration Tab

Integration of the c(*s*) distribution is typically performed in a separate software; however, BASIS allows for seamless connection between analysis in SEDFIT to an integration platform. The integration function in BASIS allows precise integration via an interpolation of data between data points in the c(*s*) distribution. The Integration Tab shows an interactive display of the c(*s*) distribution, where the user may zoom and pan around the distribution, as well as hover the cursor over the plot to more precisely determine the sedimentation coefficient ranges desired for the integration. Ranges can also be excluded from the analytical response region. This is especially useful for ignoring the half-peak at 0 S, as these half-peaks are typically caused by baseline offsets in the data, not sedimenting particles of interest ^10^. For AAV applications, it is sometimes desirable to exclude species sedimenting slower than the empty capsid and species sedimenting faster than the full capsid, such that the % areas calculated directly reflect on the ratio of capsid loading state. When integration ranges are entered and the integration is initiated, the plotted c(*s*) distribution becomes filled for easy visualization of the ranges and areas. A table also populates to show total peak areas (measured absorbance or fringes), % areas, and signal-average sedimentation coefficients. Lastly, the integration results are exported to a protected output file, alongside additional information corresponding to the fit and the location of files generated during the analysis. The output generated from BASIS is encrypted and tracks the changes made to the files, thereby adding an additional layer of security for data integrity purposes.

### 5. Analyst: QC Results Tab

Once one or more results have been exported by the analyst, the results can be read into the QC Results Tab. This tab allows multiple results to be compared, such as a series of replicates. The c(s) distributions can be viewed in the same plot, and a table is generated showing the average, standard deviation, and coefficient of variation for each specified range stemming from the previously performed integrations. Next, a PDF report may be generated with a 21 CFR Part 11-compliant electronic signature from the analyst. This report contains all information from the analysis procedure, including fit information, directories of scans used in the analysis, individual integrations, the c(*s*) overlay, the complete table with statistics on the integrated regions, time stamps for analysis/experiment, and electronic signatures.

### 6. Reviewer: Review Results

At this point, the analyst has completed their analysis and applied their signature to the report. Next, a separate user may login to BASIS under the role of Reviewer (on the same or a separate computer). The reviewer then is able to load in individual results generated by the analyst, and they are presented with the same Data Tab, Visualization Tab, and Integration Tab (read-only) that the analyst observed.

### 7. Reviewer: QC Review

After careful inspection of the individual analyses, the reviewer may load all relevant results into the QC Review Tab. This tab presents a similar view to the QC Results tab – the c(*s*) overlay and table of results are shown. The reviewer may then approve or reject the results by applying their own 21 CFR Part 11-compliant electronic signature to the PDF report. BASIS exhibits a feature which allows the Reviewer to perform these tasks from a separate device as the Analyst rather than necessitating the use of a single device for all work done throughout the workflow. If there were duplicate or triplicate data analyzed by the Analyst, BASIS checks the output files ensuring that exact same set of data files are used during the review process.

### 8. Features Implemented for 21 CFR Part 11 Compliance

The regulations set in the 21 CFR Part 11 guidance were considered during the design of BASIS and are explicitly met by the software. Examples of these features include an Administrator User account that can manage other users and make global changes to the behavior of the program. Detailed electronic records are generated at every step of data analysis and maintained in a read-only Audit Trail. The final electronic record that contains the results and parameters used for analysis is time-stamped and electronically signed by an Analyst and Reviewer. There is redundancy of data analysis parameters, such that sufficient information to reproduce an analysis can be obtained from multiple locations. Furthermore, once an analysis is generated, the analysis or files contained within cannot be altered. If a result is modified, BASIS produces an error and prohibits further use of that result.

### 9. Added Features and Improvements over the stand-alone SEDFIT

The BASIS platform also allows for improvement of the user workflow. Examples include pre-saved Parameter Sets, the Meniscus Finder, automatic saving of screenshots and distribution files, creating overlays, integration with optional exclusion ranges, as well as being able to easily save analyses for later reference.

A research-grade implementation of BASIS will also be made available. This will remove the components necessary for cGMP compliance, such as entering passwords. The standard BASIS installation also provides a way of switching between cGMP and User Analysis modes, such that BASIS can be quickly used for informal analyses or non-cGMP work.

BASIS currently reads absorbance and interference data. Intensity data must first be converted to pseudo-absorbance before analysis ^26,30,31^. We prefer to collect cGMP data in absorbance mode to avoid additional data handling and processing prior to analysis.

### 10. Application to the Optima AUC

The newer Optima AUC produces data of a format consistent with the earlier models. As such, BASIS is compatible with and intended for use with data from the Optima AUC.

### 11. Computer Software Validation (CSV)

Per the 21 CFR Part 11 regulations, software used in the processing and analysis of data collected for cGMP usage must undergo a computer software validation (CSV). A complete CSV has been performed with BASIS to ensure the software is secure, accurate, and fit for the intended purpose.

## Results

To demonstrate the usage of BASIS for cGMP-compliant analysis, triplicate analyses of Bovine serum albumin (BSA), NISTmAb reference material, and simulated AAV datasets were performed. The workflow described above was used within BASIS. Briefly, the directory was chosen in BASIS, which initiated the Meniscus Finder and presented a preview of the data with the meniscus, meniscus limits, and left data fit limit automatically set. No adjustments to the meniscus position or limits were necessary – the algorithm is highly accurate and robust (Figure 3). A custom parameter set was saved for the analysis of BSA, whereas the mAb and AAV presets were chosen for those respective analyses. SEDFIT was launched from within BASIS once the appropriate parameter set was selected. Integration was performed for each individual analysis and results saved. Next, the triplicates were loaded into the QC Results Tab to obtain the c(*s*) overlay and integration statistics, and a report was generated. The c(*s*) overlay for the BSA, NISTmAb, and AAV triplicate analyses are shown in Figure 4. Tables 1 – 3 list the integration statistics for each set of analyses. In addition to the weight-averaged sedimentation coefficient (S) determined for each region and the respective quantitation, the BASIS output allows the user to calculate a total area for the analytical response region. The latter parameter is important for analytical methods that include a requirement for concentration or total signal.

**Table 1.**
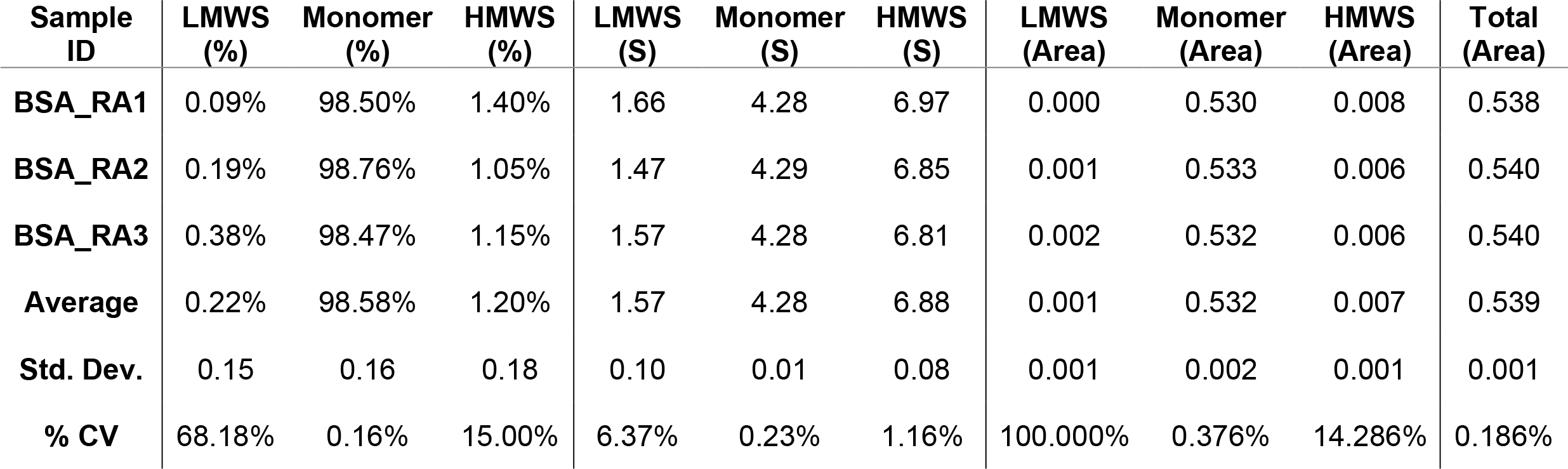
BASIS-reported integration of the BSA triplicate analysis.

| Sample ID | LMWS (%) | Monomer (%) | HMWS (%) | LMWS (S) | Monomer (S) | HMWS (S) | LMWS (Area) | Monomer (Area) | HMWS (Area) | Total (Area) |
| --- | --- | --- | --- | --- | --- | --- | --- | --- | --- | --- |
| BSA_RA1 | 0.09% | 98.50% | 1.40% | 1.66 | 4.28 | 6.97 | 0.000 | 0.530 | 0.008 | 0.538 |
| BSA_RA2 | 0.19% | 98.76% | 1.05% | 1.47 | 4.29 | 6.85 | 0.001 | 0.533 | 0.006 | 0.540 |
| BSA_RA3 | 0.38% | 98.47% | 1.15% | 1.57 | 4.28 | 6.81 | 0.002 | 0.532 | 0.006 | 0.540 |
| Average | 0.22% | 98.58% | 1.20% | 1.57 | 4.28 | 6.88 | 0.001 | 0.532 | 0.007 | 0.539 |
| Std. Dev. | 0.15 | 0.16 | 0.18 | 0.10 | 0.01 | 0.08 | 0.001 | 0.002 | 0.001 | 0.001 |
| % CV | 68.18% | 0.16% | 15.00% | 6.37% | 0.23% | 1.16% | 100.000% | 0.376% | 14.286% | 0.186% |

**Table 2.**
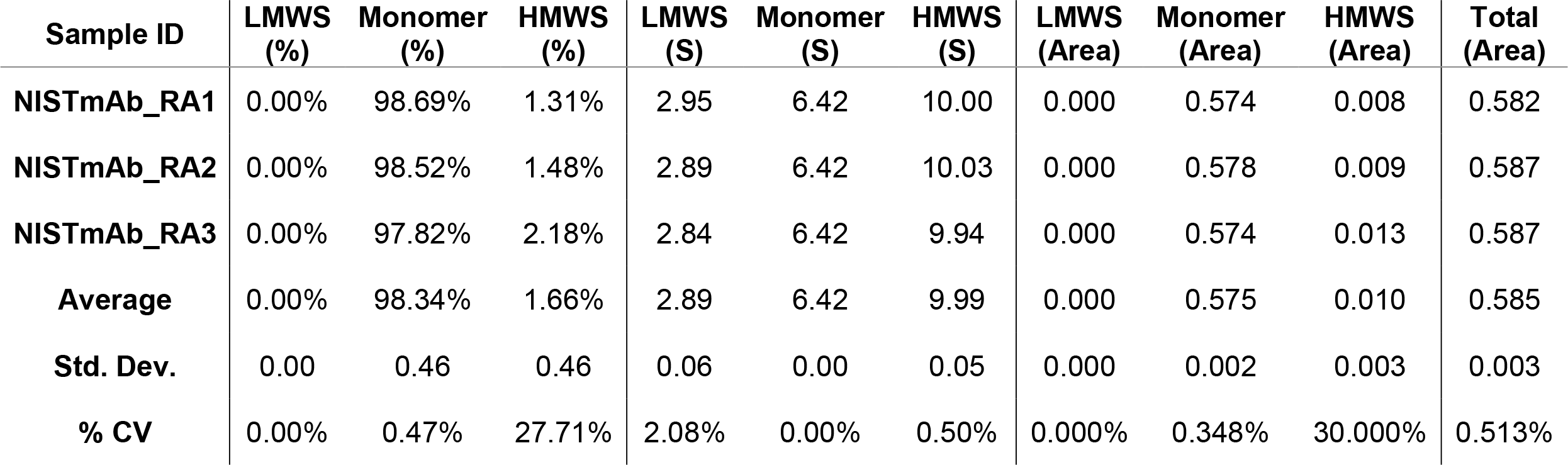
BASIS-reported integration of the NISTmAb triplicate analysis.

| Sample ID | LMWS (%) | Monomer (%) | HMWS (%) | LMWS (S) | Monomer (S) | HMWS (S) | LMWS (Area) | Monomer (Area) | HMWS (Area) | Total (Area) |
| --- | --- | --- | --- | --- | --- | --- | --- | --- | --- | --- |
| NISTmAb_RA1 | 0.00% | 98.69% | 1.31% | 2.95 | 6.42 | 10.00 | 0.000 | 0.574 | 0.008 | 0.582 |
| NISTmAb_RA2 | 0.00% | 98.52% | 1.48% | 2.89 | 6.42 | 10.03 | 0.000 | 0.578 | 0.009 | 0.587 |
| NISTmAb_RA3 | 0.00% | 97.82% | 2.18% | 2.84 | 6.42 | 9.94 | 0.000 | 0.574 | 0.013 | 0.587 |
| Average | 0.00% | 98.34% | 1.66% | 2.89 | 6.42 | 9.99 | 0.000 | 0.575 | 0.010 | 0.585 |
| Std. Dev. | 0.00 | 0.46 | 0.46 | 0.06 | 0.00 | 0.05 | 0.000 | 0.002 | 0.003 | 0.003 |
| % CV | 0.00% | 0.47% | 27.71% | 2.08% | 0.00% | 0.50% | 0.000% | 0.348% | 30.000% | 0.513% |

**Table 3.** BASIS-reported integration of the AAV triplicate analysis.

| Sample ID | LOC (%) | Empty Capsid (%) | Partial Capsids (%) | Full Capsid (%) | HOC (%) | LOC (S) | Empty Capsid (S) | Partial Capsids (S) | Full Capsid (S) | HOC (S) |
| --- | --- | --- | --- | --- | --- | --- | --- | --- | --- | --- |
| AAV_RA1 | 2.98% | 26.02% | 14.47% | 52.64% | 3.88% | 14.3 | 64.4 | 80.1 | 95.9 | 116.0 |
| AAV_RA2 | 2.98% | 25.96% | 14.66% | 52.44% | 3.97% | 14.3 | 64.4 | 80.1 | 95.9 | 116.3 |
| AAV_RA3 | 3.05% | 26.14% | 14.63% | 52.36% | 3.82% | 14.2 | 64.4 | 80.3 | 96 | 116.2 |
| Average | 3.00% | 26.04% | 14.59% | 52.48% | 3.89% | 14.3 | 64.4 | 80.2 | 95.9 | 116.2 |
| Std. Dev. | 0.04 | 0.09 | 0.1 | 0.14 | 0.08 | 0.1 | 0.0 | 0.1 | 0.1 | 0.2 |
| % CV | 1.33% | 0.35% | 0.69% | 0.27% | 2.06% | 0.70% | 0.00% | 0.10% | 0.10% | 0.20% |

| Sample ID | LOC (Area) | Empty Capsid (Area) | Partial Capsids (Area) | Full Capsid (Area) | HOC (Area) | Total (Area) |
| --- | --- | --- | --- | --- | --- | --- |
| AAV_RA1 | 0.024 | 0.209 | 0.116 | 0.423 | 0.031 | 0.803 |
| AAV_RA2 | 0.024 | 0.209 | 0.118 | 0.421 | 0.032 | 0.804 |
| AAV_RA3 | 0.024 | 0.210 | 0.118 | 0.421 | 0.031 | 0.804 |
| Average | 0.024 | 0.209 | 0.117 | 0.422 | 0.031 | 0.804 |
| Std. Dev. | 0.000 | 0.001 | 0.001 | 0.001 | 0.001 | 0.001 |
| % CV | 0.000% | 0.478% | 0.855% | 0.237% | 3.226% | 0.124% |

**Fig. 3.**
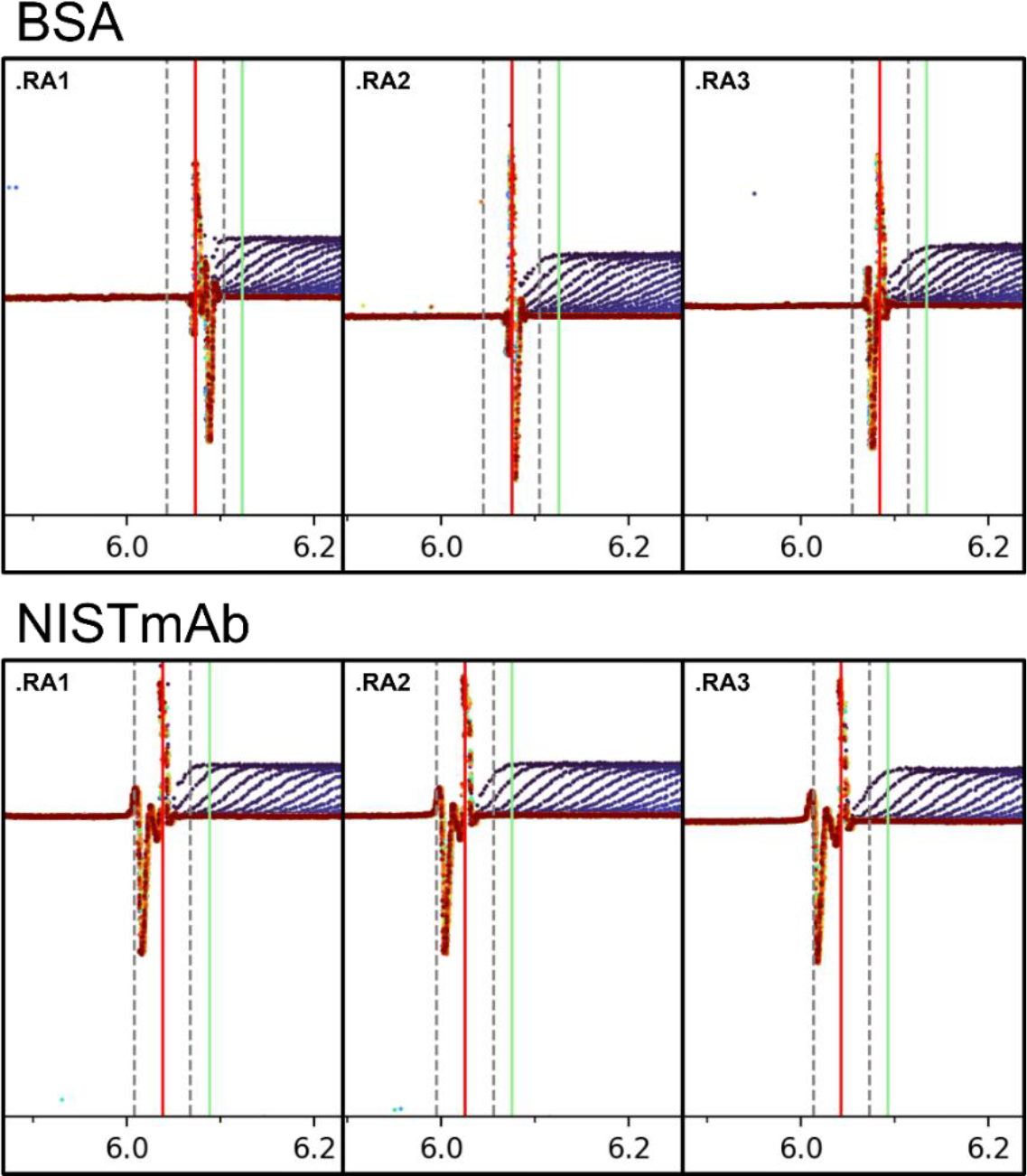
Screenshots of a portion of the Meniscus Finder window showing the accuracy of the algorithm. Raw data are shown as colored symbols ranging from dark blue to red, corresponding to the time of the scan. The x-axis displays the radius (cm), while the y-axis displays absorbance (AU). The meniscus position (solid red line), meniscus fit limits (dashed grey lines), and data fit limit (solid green line) were automatically set by the Meniscus Finder to the values shown in the screenshots.

**Fig. 4.**
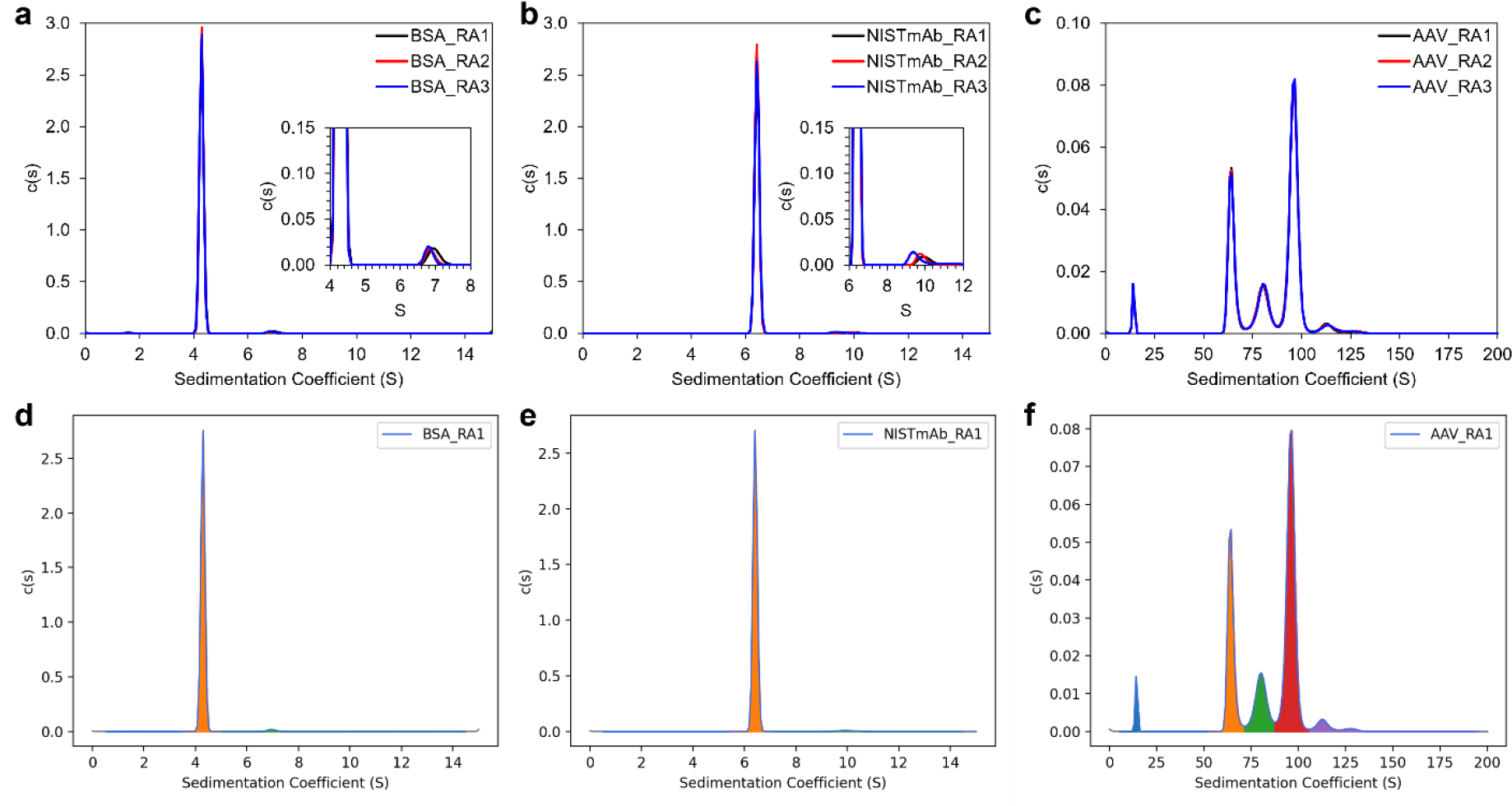
The c(s) overlays from the (**a**) BSA, (**b**) NISTmAb, and (**c**) AAV triplicate analysis using BASIS. The insets in (**a**) and (**b**) show the minor species observed. The integrated regions are shown in panels (**d**) – (**f**).

## Discussion

Analytical ultracentrifugation (AUC) is an extremely versatile and rigorous technique that is well-suited for the characterization of pharmaceuticals. It has long been used for antibody trace aggregate quantitation ^5,10^ and more recently been proven to be the “gold standard” approach for the quantitation of AAV capsid payload, as it can resolve empty, partial, and full capsids based on mass and density ^18–20,23^. While HPLC-based techniques such as SEC-MALS and ion exchange chromatography have seen cGMP analysis software in place for years now, AUC data collection and analysis software were left behind. This manuscript presented a discussion and implementation of a cGMP workflow for SV-AUC via a newly developed program – BASIS, that interfaces with the commonly used SEDFIT analysis software.

BASIS includes both administrator and user roles, electronic signatures and timestamps, data integrity checks, and an audit trail. These features are implemented to meet the 21 CFR Part 11 regulatory requirements. Other features include the Meniscus Finder, input parameter presets, integration with a specified analytical response region, and report generation with overlays and statistics tables, all of which simplify the analysis process. Another important feature is the ability of a Reviewer to review (i.e., for scientific or QC purposes) the analysis and approve or reject the report generated by the Analyst.

With the recent release of SEDFIT v16.50 came the ability to control SEDFIT from external sources (command line and MATLAB) ^32^. BASIS makes use of this functionality and provides a single and comprehensive program that covers the data analysis steps with audit trails, documentation of the experimental and data analysis procedures, timestamps, encryption, maintenance of data integrity and ultimately 21 CFR part 11 compliance. With the introduction of BASIS, the long-held belief that AUC is primarily for R&D characterization but not for the cGMP/QC environment is dispelled. With BASIS, AUC experiments can be implemented in the cGMP/QC environment in a robust manner that meets the 21 CFR Part 11 compliance standards.

## Materials and Methods

### SV-AUC data collection

Bovine serum albumin (BSA; Reference Material^®^ 927f) and monoclonal antibody (NISTmAb; Reference Material^®^ 8671) were obtained from the National Institute of Standards and Technology (NIST). Solutions were diluted with PBS (Gibco 1xPBS, pH 7.4; Fisher Scientific) to approximately 0.6 OD at 280 nm in a 1 cm pathlength quartz cuvette, as measured by a Perkin Elmer LAMBDA 365+ double-beam UV/Vis spectrometer. Data were collected on triplicate samples within the same experiment.

SV-AUC data were collected at 50,000 rpm using a 4-hole rotor on a Beckman Coulter Optima XL-A. An absorbance wavelength of 280 nm was used. A standard 2-sector, 1.2 cm pathlength Epon centerpiece was assembled with quartz windows. All samples were allowed to achieve thermal equilibrium (20 °C) over 1.5 hours under vacuum at 0 microns.

Simulation of the AAV dataset was performed using SViMULATE version 1.0.6 ^33^ was previously described ^20^. The scanning frequency was changed to 180 s to reflect the typical time required to scan all three cells on an XL-A/XL-I instrument. The scanning frequency of an Optima XL-A AUC is approximately 60 s for three cells. To generate triplicate datasets, the simulation was performed three times so as to generate new and unique random noise profiles across the three datasets.

### BASIS Meniscus Finder

The meniscus fitting limits and left fit limit were automatically set based on the calculated meniscus position. The meniscus position was floated during fitting to the c(*s*) model.

### Sedimentation velocity analysis

Data were analyzed using the c(*s*) model ^27^ in SEDFIT version 16.50 ^27,32^. The continuous c(*s*) distribution analysis assumes ideal, non-interacting species. The meniscus position and frictional ratio (*f*/*f*_0_) were fitted during the analysis. Maximum entropy regularization was applied, with a confidence level of 0.95 for the BSA and NISTmAb datasets or 0.68 for the AAV dataset.

## Statements and Declarations

### Competing Interests

All authors are employees or consultants of BioAnalysis, LLC.

### Funding Statement

This work was funded by BioAnalysis, LLC.

## Acknowledgements

The authors thank Dr. Peter Schuck for assistance on the external control of SEDFIT, Dr. David Hayes for initial assistance with BASIS, Dr. John Philo for discussions around his suite of programs, and Dr. Borries Demeler for discussions around cGMP ULTRASCAN.

## Data Availability

The simulated AAV dataset, as well as the experimental BSA and mAb datasets are available from the authors upon reasonable request.

## Author Contributions

AEY performed formal analysis and wrote the manuscript

ESG wrote BASIS

VZ-R performed analysis

NF and HMC collected data

LNP conceived the project

AEY, ESG, MTD, and LNP contributed ideas to the design and functionality of BASIS

All authors reviewed the manuscript

## Abbreviations

21 CFR: Title 21 of the Code of Federal Regulations
AAV: Adeno-associated virus
AUC: Analytical ultracentrifugation
BSA: Bovine serum albumin
cGMP: Current Good Manufacturing Practices
mAb: monoclonal antibody
SOP: Standard Operating Procedure
SV-AUC: Sedimentation velocity analytical ultracentrifugation

## Notes

### Competing Interest Statement

All authors are employees or consultants of BioAnalysis, LLC. This work was funded by BioAnalysis, LLC.

